# Regression from pathological hypertrophy is sexually dimorphic and stimulus-specific

**DOI:** 10.1101/678540

**Authors:** Deanna Muehleman, Alison R. Swearingen, Leslie A. Leinwand

## Abstract

**Aims:** Pathological cardiac hypertrophy is the result of increased cardiomyocyte size, leading to thickening of the left ventricular walls and a decrease in the left ventricular chamber. With early treatment of the underlying cause, cardiac hypertrophy can be reversed in some individuals, while it persists in others. Here, we investigate mechanisms leading to regression of pathological cardiac hypertrophy in two mouse models, in addition to defining the sex differences associated with hypertrophy and regression.

**Methods and Results:** Two pathological hypertrophic stimuli were used in male and female mice (Isoproterenol or Angiotensin II). The stimulus was removed after 7 days of treatment, then the left ventricle was studied at intervals up to 7 days following the removal of the stimulus. Following Isoproterenol removal, male hearts returned to baseline sizes in 4 days while it took 7 days for female hearts to regress. After Angiotensin II removal, the left ventricular masses of males and females did not regress. ERK1/2 was activated in response to both Isoproterenol and Angiotensin II in males, then decreased back to baseline one day after stimulus removal. Expression of ECM genes was greater in response to Angiotensin II and remained elevated longer after Angiotensin II removal, compared to Isoproterenol. Further, collagen content may be playing a role in the irreversible state of Angiotensin II induced hypertrophy as hydroxyproline content was increased following the removal of Angiotensin II in both males and females.

**Conclusions:** Regression of pathological cardiac hypertrophy is possible in some people and in some mouse models; however, the ability for the heart to regress is dependent on the stimulus and biological sex. Further, molecular changes including cellular signaling, protein degradation pathways and the formation of a fibrotic network may contribute to the ability to reverse pathological cardiac hypertrophy and are stimulus- and sex-dependent.

**Translational Perspective:** Pathological cardiac hypertrophy is a major risk factor for mortality. If cardiac hypertrophy persists for an extended time, there can be many maladaptive changes to the myocardium. With early treatment of the underlying cause, cardiac hypertrophy can be reversed in some individuals, but not in others. While cardiac hypertrophy has been studied extensively, very little is understood about regression of cardiac hypertrophy. It is important that we have a better understanding of mechanisms leading to regression and why this process might not be reversible in some individuals.

## 1. Introduction

Cardiac hypertrophy is a major risk factor for mortality and can be caused by high blood pressure, diabetes, mutations in sarcomeric proteins, and aortic valve stenosis ^(1–4)^. In mammals, this hypertrophy is the result of increased cardiomyocyte size, leading to thickening of the left ventricular (LV) walls and a decrease in the LV chamber. If cardiac hypertrophy persists for an extended time, there can be many maladaptive changes to the myocardium including: cell death, fibrosis and lengthening and thinning of the cardiomyocytes, ultimately leading to cardiac dilation and potentially, heart failure^(5)^. In early stages of cardiac hypertrophy, the increase in cardiomyocyte size is compensatory to normalize ventricular wall pressure; with early treatment of the underlying cause, cardiac hypertrophy can be reversed. However, there are varying degrees of regression of LV hypertrophy in hypertensive patients based on the treatment they receive^(6)^. Angiotensin II (Ang II) receptor antagonists, calcium channel antagonists and angiotensin converting enzyme inhibitors significantly decreased LV mass index and induced decreases of 13%, 11% and 10%, respectively^(6)^. However, diuretics and β-adrenergic receptor blockers did not affect LV mass index^(6)^. In addition, weight loss and decreased sodium intake led to similar decreases in LV mass index as patients that received anti-hypertensive drug treatments ^(7)^. Following aortic valve replacement for aortic stenosis, patients experienced LV mass reductions ranging from 17%-31%, but these changes were strongly associated with the severity of LV hypertrophy before surgery^(4)^. Overall, the longer the heart experienced a pathological stress and the increased work load on the heart persisted, the more maladaptive changes occurred, and hypertrophy was less likely to be reversible ^(5, 8)^. There are notable cases in which regression of chronic pathological cardiac hypertrophy occurred more readily, including bariatric surgery in which significant decreases in LV mass ranged from 21%-30%^(9–12)^ or a Left Ventricular Assist Device (Dores, #6551) in which significant decreases in LV mass ranged from 28%-41%^(13, 14)^. While many mechanisms that lead to cardiac hypertrophy are known, very little is understood about regression of cardiac hypertrophy.

Models for regression of cardiac hypertrophy include a de-banding from transverse aortic constriction (TAC)^(15–19)^, or the removal of a hypertrophic agonist such as Isoproterenol (Iso) or Ang II^(20–22)^. Each of these models demonstrated regression after approximately 7 days, and there was a return to baseline of pathologic gene expression markers such as ANF ^(18, 20, 22)^, BNP ^(17)^, COL1A1^(17)^ and a normalization of the functional response such as ejection fraction ^(17, 19)^ and cardiac output^(19, 21)^. However, each of these studies only reported a single timepoint after which regression was already complete. In addition, while one group reported a possible state of irreversible hypertrophy with one week of de-banding after chronic TAC (6 weeks), there was still a reversal of pathological gene expression including, MYH7 and ACTA-1^(17)^. However, the rate of regression, and the many molecular mechanisms that occur during the process of regression, are still not known. A better understanding of the immediate responses and the progression of regression, along with a model of incomplete hypertrophy regression, will be important to understand why some patients experience regression and others do not and sex differences have not been adequately addressed.

Sex Differences in cardiac diseases have been observed for many years now, with some differences observed in the rates and extent of hypertrophy and/or regression. For example, with aortic stenosis, although females experienced cardiac hypertrophy more often, and to a greater extent, females also regressed from hypertrophy faster than males ^(23)^. In addition, males had higher expression of collagen I & III and matrix metalloproteinase 2 genes due to aortic stenosis, which may be a factor contributing to the slower rates of regression in males ^(23, 24)^. In the studies of LVAD placement, while a few of the patients receiving an LVAD were female, most were males ^(13, 14, 25)^. Although sex was accounted for in the demographics, sexes were combined when analyzing LV mass differences and molecular changes. Similarly, patients receiving bariatric surgery were primarily female, but sexes were combined when analyzing results. However, there was one study that showed male sex is independently associated with an increase in the 1-year mortality rate post-bariatric surgery^(26)^. In the rodent models of regression of hypertrophy, de-banding from TAC or removal of the hypertrophic agonist (Iso or Ang II), were carried out with either only male rodents ^(16, 17, 20–22)^, the sex was not stated ^(18)^, or the results of the sexes were combined ^(15)^. We are just beginning to fully understand the many molecular mechanisms that underlie sex differences, even at baseline^(27)^. Therefore, it is essential to define sexually dimorphic cardiac differences, both at baseline, and in response to stress.

Here, we compare cardiac hypertrophy and regression induced by two different pathological hypertrophic stimuli, Iso or Ang II in male and female mice. In addition, we compare the sexes at baseline for many cardiac parameters and find significant differences. Although both agonists caused hypertrophy, as expected, the extent of regression of hypertrophy was distinct between the sexes and the agonist. There were differences in activated signaling pathways, autophagy and proteasome activity, and extracellular matrix composition depending on the hypertrophic stimulus and/or the biological sex.

## 2. Methods

### 2.1 Animals and treatments

All animal treatments were approved by the Institutional Animal Care and Use Committee at the University of Colorado Boulder (Protocol #2351) and are in accord with the NIH guidelines. Wild-type, 10-12-week-old C57Bl/6 male and female mice (Jackson Laboratories) were fed *ad libitum* standard rodent chow and housed in a 12-hour light/dark cycle. Mice were treated with Iso (30mg/kg/day) or Ang II (2.88mg/kg/day) for 7 days. Iso was prepared in 1μM Ascorbic Acid, diluted in sterile saline. Ang II was diluted in sterile saline. Iso and Ang II were released through osmotic mini-pumps (Alzet model 2001). Mice were anaesthetized with Isoflurane (3%) via spontaneous inhalation. Surgical procedures were performed on a 37°C re-circulated heating pad. The analgesic buprenorphine was used at 1mg/kg. To study regression, the osmotic pumps were removed after 7 days of Iso/Ang II; the same surgical procedure was used as placement of the mini-pump. Mice were euthanized by first anaesthetizing with Isoflurane (3%) via spontaneous inhalation, then the heart was removed. Mice were sacrificed at either 7 days of Iso/Ang II, indicating peak hypertrophy, or at Post-Stimulus Day 1, 4 & 7 (P1, P4 & P7 respectively). There were vehicle controls (1μM Ascorbic Acid in sterile saline for Iso or sterile saline alone for Ang II) at each time-point. Hearts were dissected, and the left ventricle was weighed then flash frozen in liquid nitrogen. Tissue was placed at −80C until further analysis.

### 2.2 Phospho-kinase array kit

Array analysis was performed according to manufactures protocol (R&D Systems, ARY003B). Briefly, tissue was homogenized in Lysis Buffer 6 and 500μg of lysate was added per array set and incubated overnight at 4°C. The arrays were washed, then incubated in the Detection Antibody Cocktail for 2 hours at room temperature. Streptavidin-HRP bead were incubated with the arrays then detection was performed using chemiluminescence.

### 2.3 Protein Isolation & Western Blots

Left ventricle tissue was homogenized in Urea Buffer (8M Urea, 2M Thiourea, 50mM Tris (pH 6.8), 75mM DTT, 3% SDS, 0.05% Bromophenol Blue). Protein concentration was determined using Pierce 600 Protein Assay Reagent (Thermo Scientific 22660) with the Pierce Ionic Detergent Compatibility Reagent (Thermo Scientific 22663). Proteins were run on 4-12% Bis-Tris gels and transferred to Nitrocellulose membranes. Membranes were blocked with 5% BSA in TBST for 1 hour at room temperature. Primary antibodies were incubated O/N in 5% BSA (TBST) at 4°C. Secondary antibodies were incubated for 1 hour at room temperature. Membranes were imaged using ECL reagent (Perkin-Elmer NEL104001EA). Quantification was determine using ImageQuant. All primary antibodies were purchased through Cell Signaling Technology: p-Akt (4058), p-mTOR (2974), p-GSK3β (9336), p-p38 (4511), p-ERK1/2 (9101), LC3 (2775) except α-Vinculin was purchased from Sigma (V9131).

### 2.4 Proteasome Activity Assay

Left ventricle tissue was homogenized in Proteasome buffer (50mM HEPES; 20mM KCl; 5mM MgCl_2_; 1mM DTT). Samples were centrifuge at 10,000xg for 30 minutes at 4°C. The supernatant was placed in a new tube and the protein concentration was determined using Pierce BCA Protein Assay (Thermo Scientific 23227). Each reaction contained 15μg protein in a final volume of 230μl Proteasome buffer. In addition, each sample had a complimentary reaction which contained the proteasome inhibitor MG132 (20nM). 230μl of the sample was prepared on a 96-well white flat bottom plate. 10μl of the fluorescent substrate was added; Suc-LLVY-AMC was used to measure chymotrypsin activity (18μM; Enzo Life Sciences BML-P802). The samples were kept on ice until this point. The reaction was started by placing the plate in the plate reader at 37°C and fluorescence was measured every 3 minutes for 60 minutes (excitation 360nm; emission 460nm). Proteasome activity was determined by calculating the change in fluorescence; this value was then subtracted from the change in fluorescence from the complimentary reaction containing MG132. Each sample was run in triplicates.

### 2.5 RNA isolation

LV tissue was homogenized in Tri Reagent (Molecular Research Center, TR118). Chloroform was added and incubated at room temperature for 15 minutes, then centrifuged at 12,000xg for 15 minutes at 4°C. The aqueous layer was removed and placed in a new tube. Isopropanol was added and incubated at room temperature for 15 minutes, then centrifuged at 12,000xg for 15 minutes at 4°C. The supernatant was removed, and the pellet was washed with 70% ethanol, then centrifuged at 7,500xg for 5 minutes at 4°C. The supernatant was removed, and the RNA was resuspended in water.

### 2.6 cDNA preparation and Quantitative Real-Time PCR

RNA was reverse transcribed using SuperScript III Reverse Transcriptase (Invitrogen 18080044) and the protocol was followed according to the manufacture. cDNA was diluted to 1μg/ul in water. Each QPCR reaction contained, 4μg cDNA + SYBR Green PCR Master Mix (Invitrogen 4309155) +12.5μM primer set. Thermocycler settings were determined used SYBR Green PCR Master Mix Protocol. ΔΔCt was calculated using 18S as a normalizer.

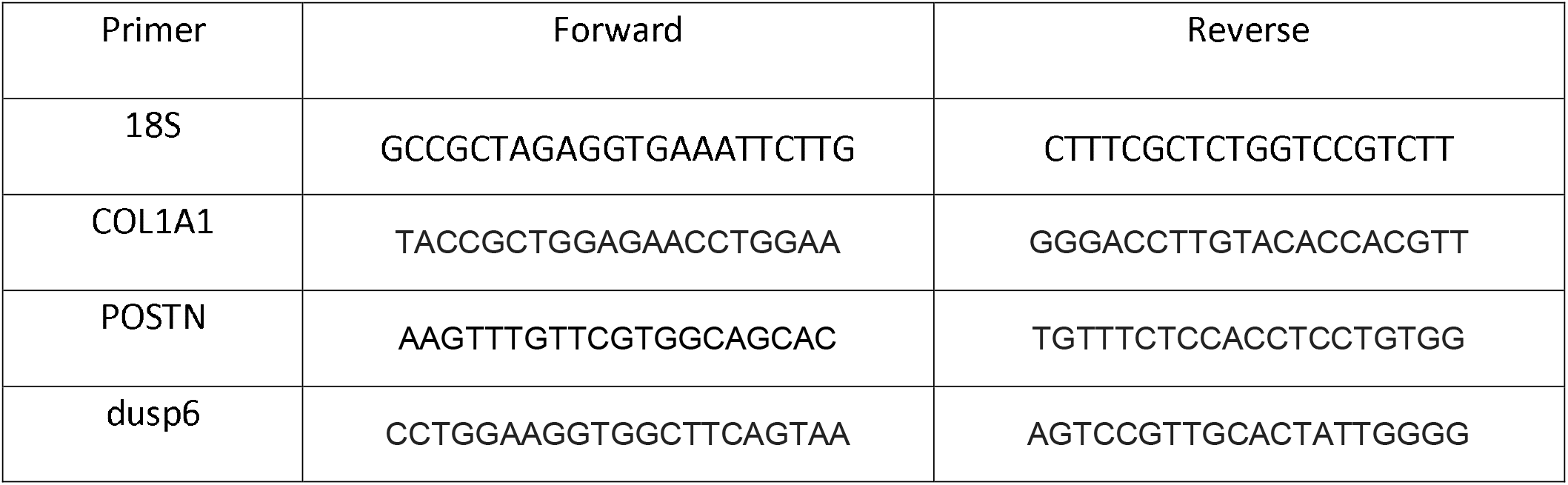

### 2.7 Hydroxyproline Assay

A hydroxyproline assay kit was used (Sigma-Aldrich; MAK008) and followed according to the manufacturer’s instructions. Briefly, 10mg of tissue was homogenized, then hydrolyzed in 12M HCl. 30μl of each sample was transferred in duplicate to a 96 well plate and allowed to dry in a 60°C oven for 60 minutes. Chloramine T/Oxidation Buffer was added to each sample and standard well, followed by the diluted DMAB Reagent. Samples were incubated for 90 minutes at 60°C. Absorbance was measured at 560nm.

### 2.8 Statistical Analysis

Statistical differences were determined between two groups using Student’s two-tailed T-test. Between multiple groups, one-way ANOVA was performed followed by Uncorrected Fisher’s LSD post-hoc test. Outliers were determined using the Grubbs test with an α<0.05. P-values of less than 0.05 were considered significant.

## 3. Results

### 3.1 Pathological cardiac hypertrophy and regression

To induce pathological cardiac hypertrophy, we used the β-adrenergic agonist, Iso, or an activator of the Renin-Aldosterone-Angiotensin-System (RAAS) (Ang II). Iso (30mg/kg/day) induced a 33.8% increase in Left Ventricle/Tibia Length (LV/TL) in males (Figure 1C). Female mice showed a 28.5%increase in LV/TL (Figure 1D). Although this male-female difference was not statistically significant, this corroborates an earlier study that showed females have a more moderate hypertrophic response to Iso treatment^(28)^. After removal of Iso, male mice showed significant regression after 1 day (P1) but were still significantly larger than the vehicle control (Figure 1A, C). They showed complete regression after 4 days (P4) and were no longer different than the vehicle control. After Iso was removed, female mice showed significant, but incomplete regression immediately at P1, similar to males (Figure 1B, D). In contrast to the males, females only showed complete regression at P7.

**Figure 1:**
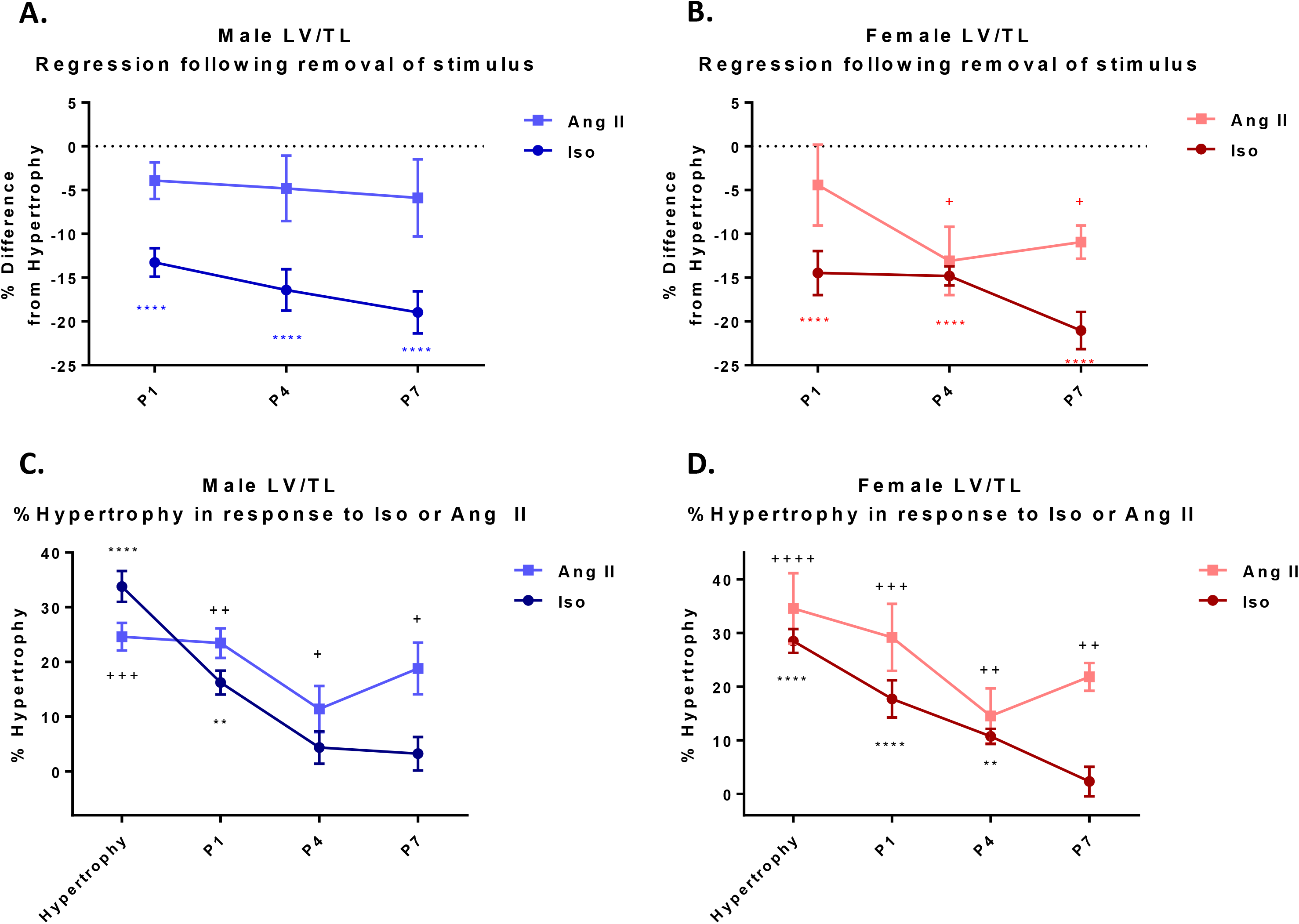
Complete regression from Iso-induced hypertrophy but not Ang II-induced hypertrophy. A. %Regression of LV/TL in males compared to hypertrophy group B. %Regression of LV/TL in females compared to hypertrophy group. C. %Hypertrophy of LV/TL in males compared to vehicle group. D. %Hypertrophy of LV/TL in females compared to vehicle group. n=6-8/group. LV/TL Left Ventricle Weight/Tibia Length; P Post Hypertrophy Day. One-way ANOVA Post hoc-Uncorrected Fisher’s LSD. */+p<.05 significance from vehicle control. * significance in Iso group. + significance in Ang II group. */+; */+ significance from hypertrophy group.

In order to determine whether a pathological response to a different stimulus would show similar patterns of hypertrophy and regression, we also used Ang II, which we hypothesized would cause irreversible hypertrophy because it has been shown to induce cardiac fibrosis in mice ^(29)^. Male mice treated with Ang II (2.88mg/kg/day), experienced an increase of 24.6% in LV/TL (Figure 1C), and female mice experienced an increase of 34.6% in LV/TL (Figure 1D), but these sex differences were not statistically significant. After Ang II was removed, males did not experience any significant regression while females experienced significant, but incomplete regression after Ang II was removed for 4 days (Figure 1A, B); however, even after 7 days, the LV weights of both males and females, remained significantly larger (approximately 20%) than the vehicle controls (Figure 1C, D). This was in stark contrast to Iso treatment, when complete regression occurred by P4 and P7 in males and females, respectively.

Therefore, this is the first report to investigate the immediate response, and the rates of regression with two different models, including one that is irreversible at the timepoints studied here, in addition to examining biological sex as a variable.

### 3.2 Up and down-regulation of the ERK1/2 pathway was common to Iso and Ang II in hypertrophy and after the removal of the agonists

We next probed which signaling pathways might be regulated during the response to agonist removal and regression of cardiac hypertrophy. We initially used a commercial phospho-protein array to quantitate the phosphorylation status of 46 signaling molecules (Supplemental Figure 1). Some of these array results were validated with Western blots. First, comparisons were made between males and females at baseline, without any treatments. Interestingly, females had significantly greater levels of active ERK1/2, Akt, GSK3β and mTOR than their male counterparts but there were no sex differences in the levels of p-p38 (Figure 2).

**Figure 2:**
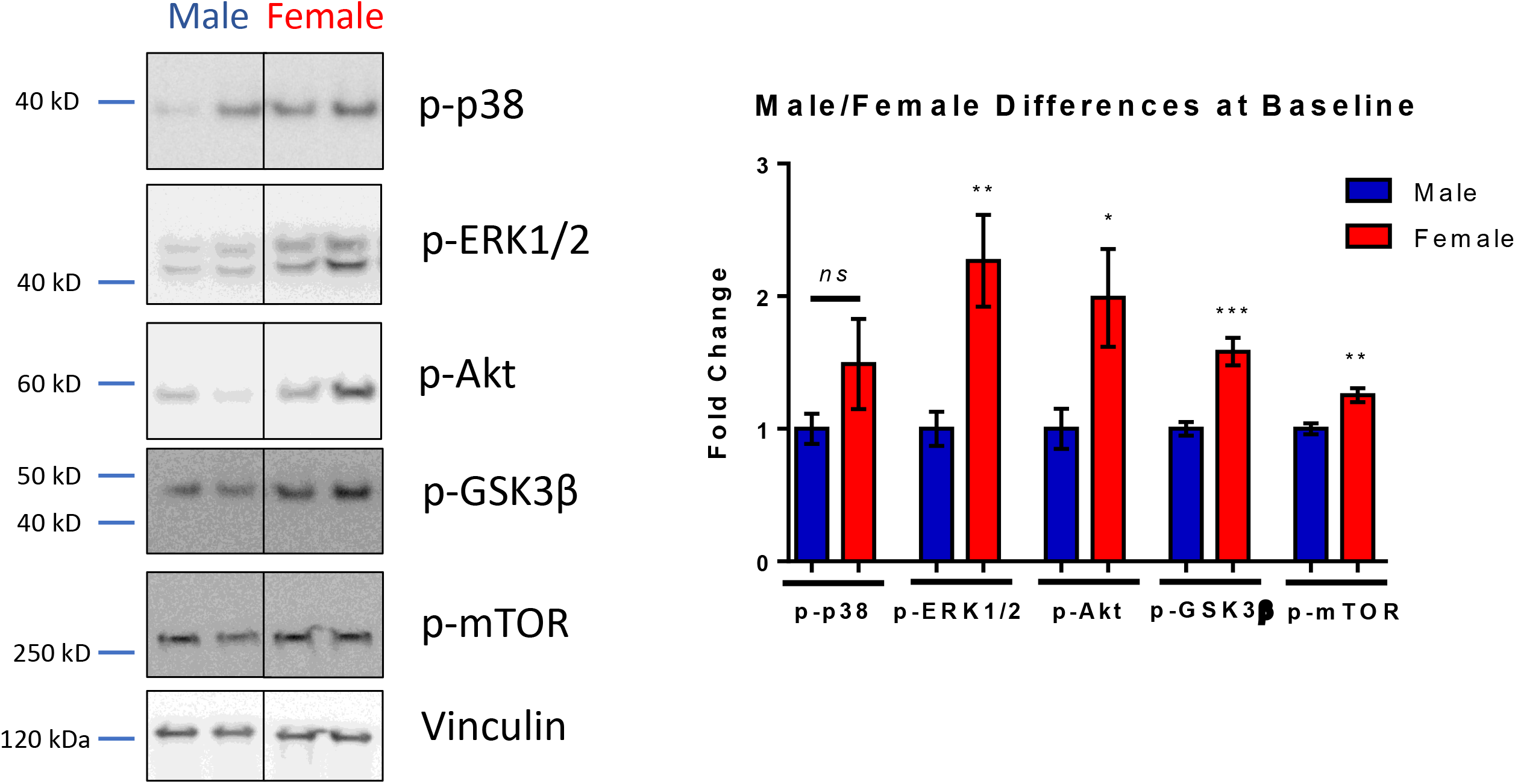
Baseline signaling activation is distinct between male and female hearts. ERK1/2, Akt, GSK3β and mTOR are more activated in females than males. There are no sex differences in active p38. Values are normalized to Vinculin. n=6-8/group. Mean ±SEM. Statistical analysis performed: Two-tailed students T-test. *p<.05.

In Iso treated male mice, p-Akt did not change significantly in response to Iso; however, it decreased during regression, compared to the hypertrophic timepoint (Supplemental Figure 2A). p-GSK3β increased in response to Iso and remained elevated at P1, then decreased at P7, compared to the hypertrophic timepoint (Supplemental Figure 2C). p-mTOR did not change in response to Iso. However, it increased during later regression at P4 and P7 (Supplemental Figure 2D). In the MAPK pathway, in males, p-p38 did not change (Supplemental Figure 2E), but p-ERK1/2 increased by 2.5-fold with Iso, then decreased to baseline levels immediately at P1 of regression (Figure 3A and Supplemental Figure 3A & B). In contrast to males, in female mice, only three kinases changed overall (p-Akt, p-GSK3β & ERK1/2) and the only changes observed were decreases during regression (Figure 3B & Supplemental Figure 2).

**Figure 3:**
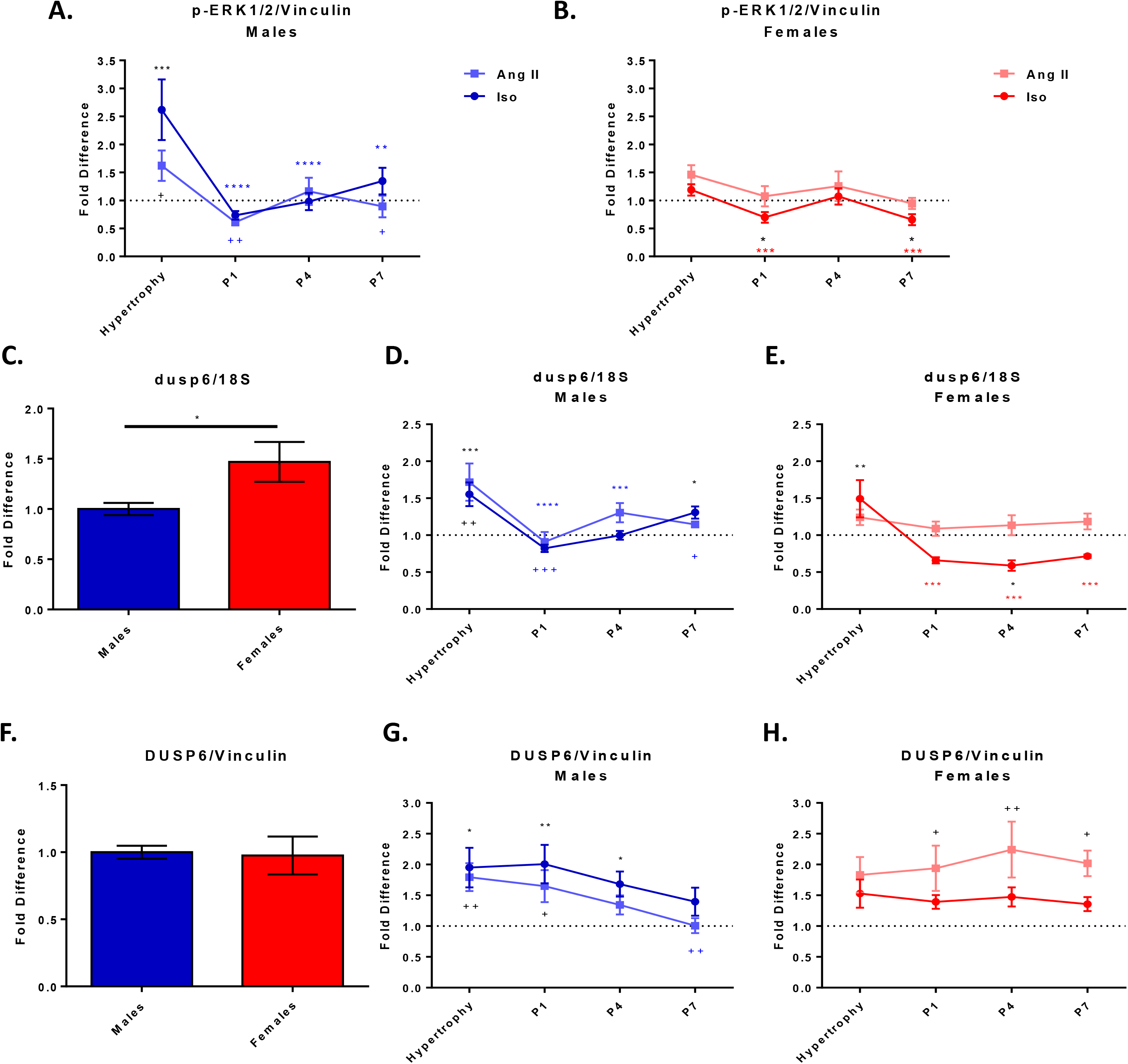
ERK1/2 activation and DUSP6 expression in response to Iso and Ang II. A. ERK1/2 is activated in male mice with Iso and Ang II which decreases after stimulus removal B. ERK1/2 is not activated in female mice with Iso or Ang II. C. dusp6 mRNA is higher in females than males at baseline. D. dusp6 mRNA increases in male mice with Iso and Ang II which decreases after stimulus removal. E. dusp6 mRNA increase in female mice with Iso, but not Ang II.F. DUSP6 protein is not differentially expressed. G. DUSP6 protein is increased in male mice with Iso and Ang II and remains elevated longer after Iso removal. H. DUSP6 protein increases after Ang II removal in female mice. Protein quantifications are normalized to Vinculin. mRNA is normalized to 18S. n=6-8/group. Mean ±SEM One-way ANOVA Post hoc-Uncorrected Fisher’s LSD. *p<.05. * significance in Iso group. + significance in Ang II group. */+significance from vehicle control. */+; */+ significance from hypertrophy.

In Ang II-treated mice, p-Akt did not change in male mice, and decreased in female mice both during hypertrophy and following the removal of Ang II (Supplemental Figure 2A & B). These results in male mice were in contrast to Iso-treated mice. We also measured the activation state of ERK1/2, as this was a predominant pathological response with Iso. As with Iso, Ang II induced an activation of ERK1/2 in males, and also consistent with Iso, levels decreased after Ang II was removed (Figure 3A and Supplemental Figure 3A & B). There were no significant changes of p-ERK1/2 in females due to Ang II (Figure 3B).

Overall, both Iso and Ang II activated Erk1/2 significantly in males and its activity decreased after removal of either agonist.

### 3.3 DUSP6/MKP3 activity may regulate p-ERK1/2 in response to Iso and Ang II hypertrophy and regression

Given that p-ERK1/2 was the most strongly activated signaling molecule during Iso- and Ang II-induced hypertrophy in males, which then decreased during regression, we hypothesized that DUSP6/MKP3 would direct the removal of the phosphate during regression. ERK1/2 is inactivated through its dephosphorylation by Dual Specificity Phosphatases (DUSPs) and DUSP6/MKP3 is highly specific for ERK1/2 ^(30)^. Previous studies showed that DUSP6 is regulated post-transcriptionally^(31)^ and post-translationally^(32)^; therefore, both mRNA and protein levels were measured. At baseline, dusp6 gene expression was significantly higher in females compared to males (Figure 3C). Iso induced a 1.5-fold increase in mRNA levels in both male and female mice (Figure 3D & E). During regression, dusp6 mRNA decreased significantly immediately after Iso removal. While these levels remained low in the females during the duration of the regression time-points, in males at P7, dusp6 increased to levels significantly higher than baseline. DUSP6 protein levels showed no differences at baseline between males and females (Figure 3F). Iso induced a nearly 2-fold increase in DUSP6 protein in males and remained significantly higher during regression until P7, when it returned to baseline levels (Figure 3G & H and Supplemental Figure 3C & D). In females, while there was a trend towards an increase in DUSP6 protein, no significant changes were observed.

Ang II resulted in a significant increase in dusp6 expression, but only in males (Figure 3D & E). This was followed by a significant reduction of dusp6 mRNA immediately following the removal of Ang II at P1 and returned to baseline. There were no significant changes observed in female mice in response to Ang II or the removal of Ang II. DUSP6 protein levels were significantly increased in male mice in response to Ang II, and while there was a trend towards an increase in females, it was not significant (Figure 3F & G and Supplemental Figure 3C & D). However, at P1, DUSP6 protein remained elevated in males and the increase in females was now significantly higher than the vehicle controls. By P4, DUSP6 returned to baseline levels in male mice. However, levels remained significantly elevated in female mice at P4 and P7.

### 3.4 Proteasome activity is differentially activated during cardiac hypertrophy and regression

During cardiac hypertrophy, there is an increase in protein synthesis ^(33)^. We hypothesized that the ubiquitin proteasome system would be involved in regression to degrade some of the proteins that accumulated during hypertrophy. Proteasome activity was determined by incubating the proteasome lysate with the proteasome chymotrypsin-like substrate, Suc-LLVY-AMC ^(34)^. First, there were no proteasome activity differences observed between males and females at baseline (Figure 4A). Next, we showed that proteasome activity increased with Iso in male mice during hypertrophy and increased further during early regression, suggesting it may be involved in protein turnover in both processes (Figure 4B). Then, at the time point when regression had occurred completely in males (P4), proteasome activity returned to baseline. In female Iso treated mice, proteasome activity remained unchanged throughout hypertrophy and regression (Figure 4C). In Ang II treated male mice, proteasome activity increased during hypertrophy, and to a greater extent compared to Iso (Figure 4B). Interestingly, the activity decreased immediately at P1. In contrast to Iso regression, proteasome activity may not be involved in the response to the removal of Ang II in males, nor in female Ang II treated mice, since no significant changes were observed (Figure 4C). The lack of proteasome activation following the removal of Ang II treated male mice, could be contributing to the irreversibility of LV weight. However, there were no differences observed between Iso or Ang II treatment in female mice.

**Figure 4:**
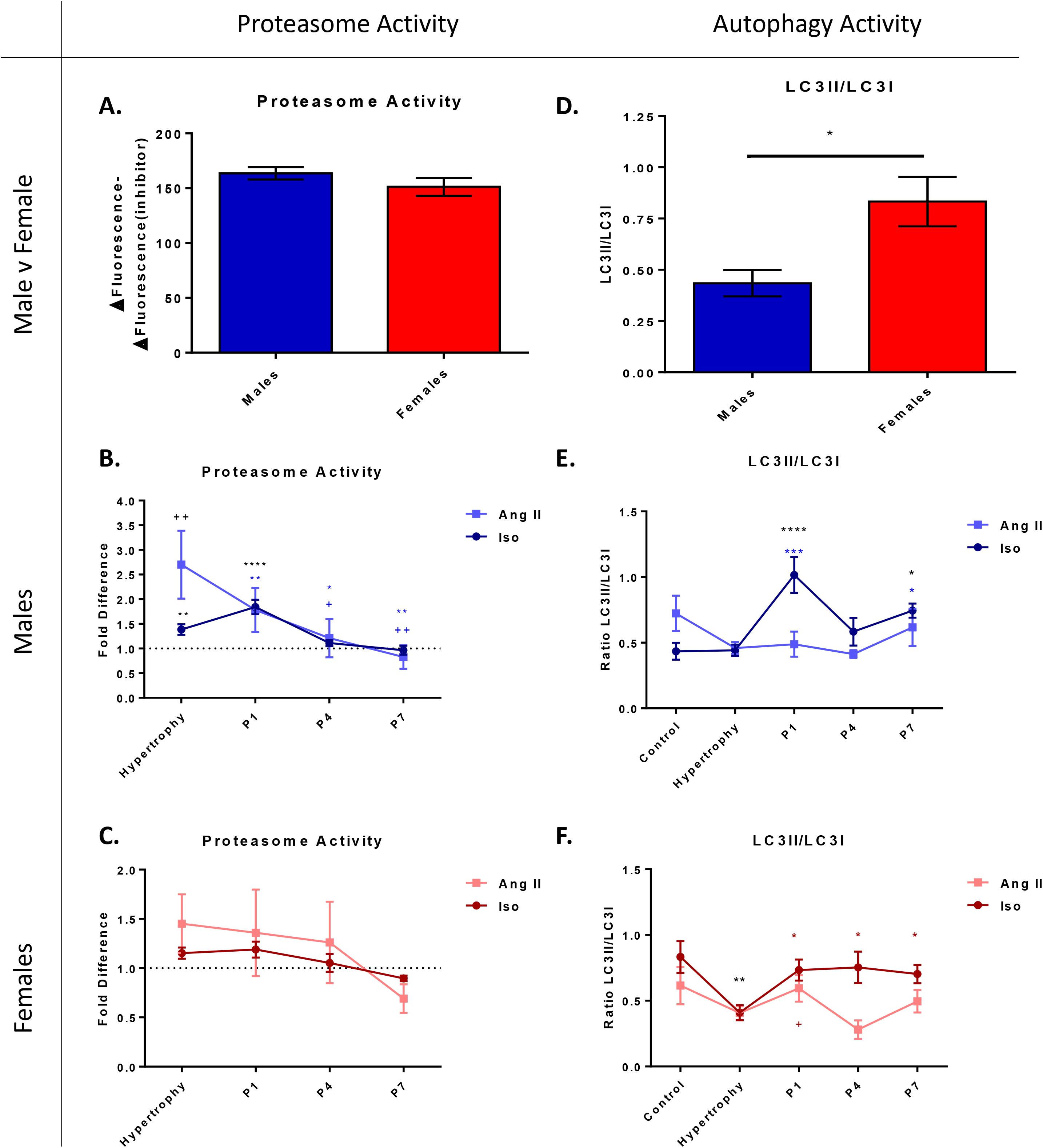
Protein degradation pathways. A. Proteasome activity is not differentially expressed. B. Proteasome activity increases in males in response to Iso and Ang II; then increases further in response to Iso removal but decreases after Ang II removal. C. Proteasome activity is unchanged in female mice. D. Autophagy activity is higher in females than males at baseline. E. Autophagy activity increases after the removal of Iso in male mice. F. Autophagy activity decreases with Iso; then increases after Iso and Ang II removal. n=6-8/group. Mean ±SEM One-way ANOVA Post hoc-Uncorrected Fisher’s LSD. *p<.05 *significance in Iso group. + significance in Ang II group. */+significance from vehicle control. */+; */+ significance from hypertrophy.

### 3.5 Autophagy is activated during regression of cardiac hypertrophy

We also hypothesized that autophagy would be activated during regression to degrade proteins that accumulated during hypertrophy. As a measure of autophagy, we quantified the protein levels of the two forms of LC3. LC3 is small protein that becomes lipidated to the growing autophagosome and we measured the amount of the lipidated form (LC3II) relative to the unlipidated form (LC3I). At baseline, females had 2-fold more LC3II/LC3I, compared to males, indicating they had higher autophagic activity (Figure 4D). Interestingly, with Iso, the amount of autophagy remained unchanged in male mice (Figure 4E) and decreased in female mice (Figure 4F). During regression, relative to the hypertrophic state, autophagy increased in both male and female mice, although the increase was greater in males. Ang II did not significantly alter the lipidation state of LC3 in males during hypertrophy or after the removal of Ang II (Figure 4E). In female mice, autophagy was unchanged with Ang II hypertrophy, but then increased at P1, and decreased by P4 (Figure 4F). These results indicate females may be attempting to promote regression via autophagy, but this increase does not have a significant influence on LV weight. Autophagy does not appear to be contributing to promoting regression from Ang II hypertrophy in male mice.

### 3.6 Extracellular matrix genes are inactivated during regression of cardiac hypertrophy

The formation of a fibrotic network in the myocardium may play a role in the ability to regress following a pathological stress. We therefore tested whether ECM genes were activated during hypertrophy and if they were inactivated during regression. We measured the expression of Collagen1a1 (COL1A1) and Periostin (POSTN) mRNAs, both of which are components of the extracellular space and contribute to the fibrotic network^(35)^. At baseline, female mice expressed significantly more COL1A1 (Figure 5A), but not significantly more POSTN (Figure 5D), compared to male mice (Figure 5A & D). During the pathological hypertrophic response induced by Iso, in both male and female mice, there was an increase in expression of both COL1A1 and POSTN (Figure 5B, C, E & F). During regression, COL1A1 expression remained elevated, then returned to baseline at P4 in both males and females (Figure 5B & C). Interestingly, there was an increase in COL1A1 expression at P7, although the reason for this is unknown. POSTN expression remained elevated in male mice at P1 and decreased to baseline levels by P4; however, it decreased to baseline in the female mice at P1 (Figure 5E, F).

**Figure 5:**
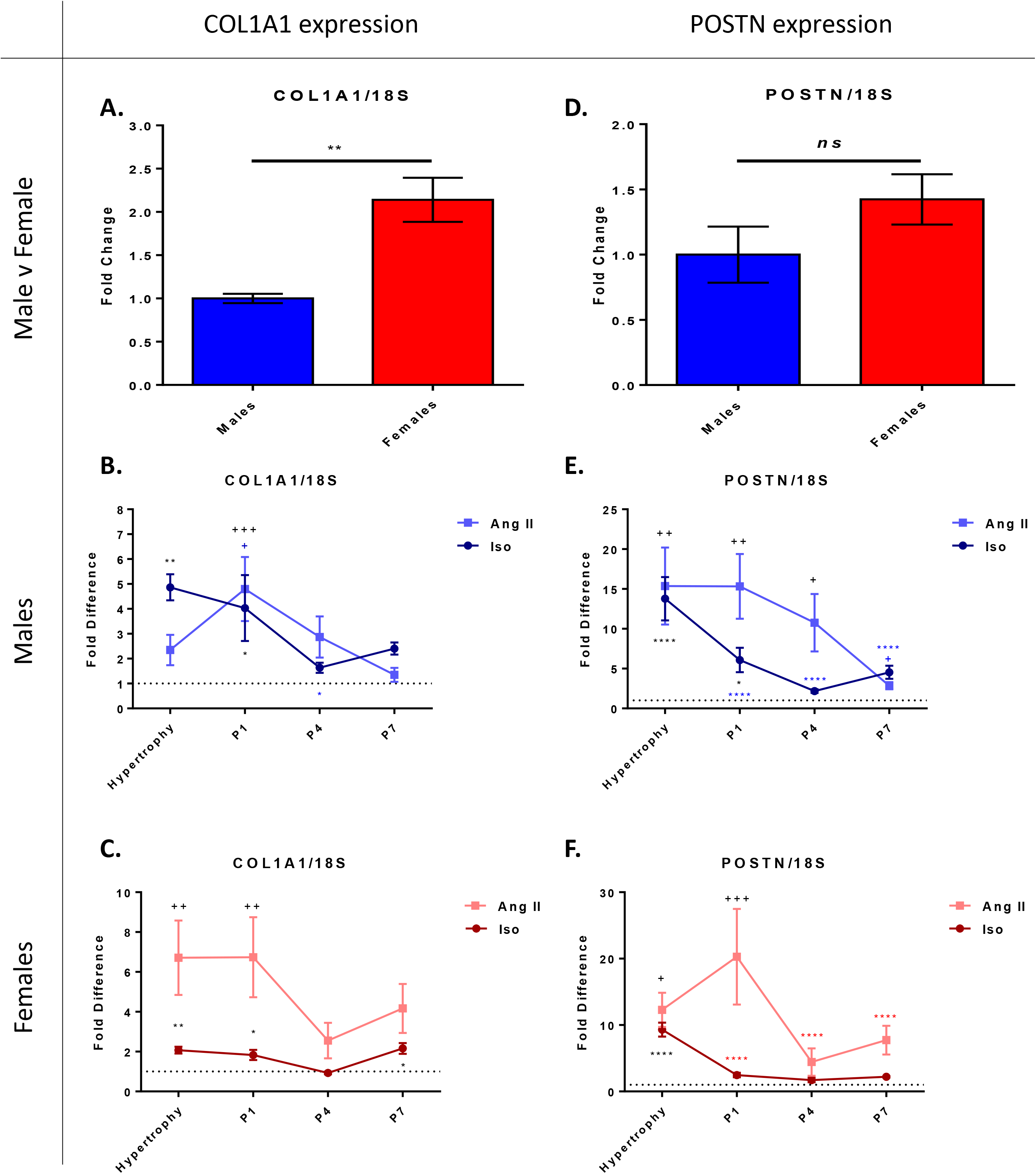
ECM gene expression has a sexually dimorphic response to Iso- and Ang II-induced hypertrophy and to the removal of the stimuli. A. COL1A1 mRNA is higher in females than males at baseline. B. COL1A1 mRNA increases in response to Iso and to the removal of Ang II in males. C. COL1A1 mRNA increases in response to Iso and Ang II followed by decreases 4 days after stimulus removal. D. POSTN mRNA is not differently expressed. E. POSTN mRNA increases in response to Iso and Ang II and decreases more slowly following the removal of Ang II compared to Iso removal. F. POSTN mRNA is increased in response to Iso and Ang II and decreases more slowly following the removal of Ang II compared to Iso. n=6-8/group. Mean ±SEM One-way ANOVA Post hoc-Uncorrected Fisher’s LSD. *p<.05 * significance in Iso group. + significance in Ang II group. */+; */+ significance from hypertrophy.

Ang II also caused increases in both COL1A1 and POSTN expression. However, Ang II induced a significantly greater response of COL1A1 in females (Figure 5C). Both agonists similarly increased POSTN mRNA with no significant sex differences (Figure 5E, F). The expression of COL1A1 had a similar trend as with Iso, and remained significantly elevated during early regression, and returned to baseline levels by P4 (Figure 5B, C). However, in contrast to the POSTN expression trend due to Iso, POSTN levels remained significantly higher following the removal of Ang II and did not regress until P7 in male mice and P4 in female mice (Figure 5E, F). Therefore, because of the higher expression levels in response to Ang II compared to Iso, and the time it took to reduce the expression levels of these ECM genes back to baseline, the formation of a fibrotic network could be contributing to the inability to regress from Ang II-induced hypertrophy.

### 3.7 Hydroxyproline content increases following the withdrawal of Ang II and did not return to baseline

Due to the increased ECM gene expression in response to Ang II, we hypothesized there would be an increase in the amount of collagen formation that is unable to be degraded and that would ultimately inhibit regression. We measured hydroxyproline content and found at baseline, females had significantly higher collagen content than males (Figure 6A). After 7 days of Ang II, there were no significant changes in either sex. However, by P1 in males, there was a significant, 1.5-fold increase, and these levels remained elevated until P7 (Figure 6B). Similar changes were seen in female mice, in that while no changes occurred immediately due to Ang II, by P4, there was a significant increase, and this increased further at P7.

**Figure 6:**
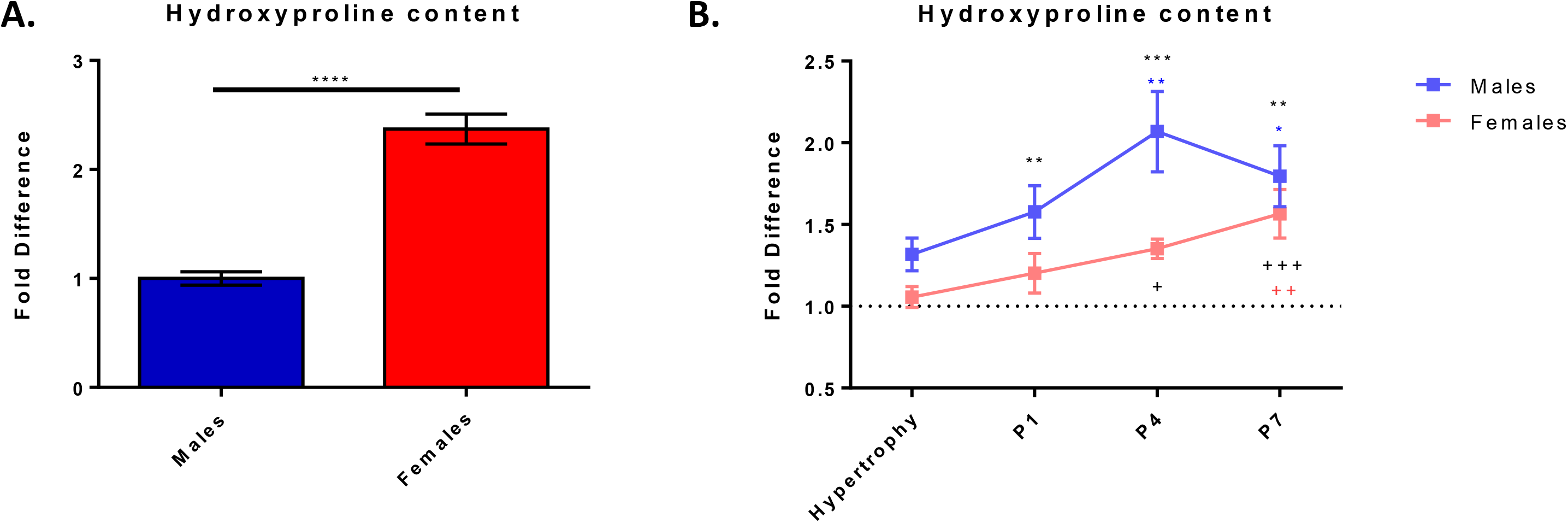
Hydroxyproline content increases following the removal of Ang II. A. Baseline hydroxyproline content is higher in female than males at baseline B. Hydroxyproline content increases following 1-day removal of Ang II in males and 4 days removal in females. n=6/group. Mean ±SEM One-way ANOVA Post hoc-Uncorrected Fisher’s LSD. *p<.05 *significance in males;. + significance in females. */+ significance from vehicle control. */+ significance from hypertrophy.

## 4. Discussion

Pathological cardiac hypertrophy can be reversed with pharmacological treatment, weight loss or surgery in some patients; however, not all patients are able to regress ^(1, 4, 6–8)^. Because cardiac hypertrophy itself is a risk factor for mortality, it is crucial that we have a better understanding of mechanisms that lead to the state of irreversibility and mechanisms that promote regression from pathological cardiac hypertrophy. To begin to understand regression, we induced hypertrophy with two different pathological agonists. Then, following the removal of the agonists, we studied the time course of regression, specific pathways that were regulated during regression, and we studied these processes in both males and females. Iso and Ang II are known to act through different pathways with Iso activating adrenergic receptors that are a class of G-protein coupled receptors on the cell surface that cause a canonical signaling cascade leading to increased concentration of Ca^2+^ in the cytosol^(36)^. This increase ultimately results in faster contractions of the cardiomyocytes and mimics the increased work load observed in the disease state. Increased adrenergic signaling via catecholamines can lead to high blood pressure and hypertrophy. Further, many people with hypertension are effectively treated with adrenergic receptor antagonists ^(37)^. Ang II, which is part of the RAAS, is also responsible for regulating blood pressure. Ang II is the main effector in RAAS, and binds to Ang II receptors (AT1R)^(38)^. Ang II induces inflammation along with many pathological hypertrophic markers (NF-κB, TNFα, MAPK & Akt signaling), and ultimately leads to an increase in blood pressure^(38)^, which leads to cardiac hypertrophy. While Iso induces cardiac hypertrophy to mimic the disease state in this mouse strain (C57Bl/6), there are reports of less activation of pathological fetal genes ^(39)^ and little evidence of fibrosis ^(40, 41)^ compared to other mouse strains. In contrast, with Ang II, there has been evidence for pathological signaling and significant cardiac fibrosis ^(29, 38)^ in the C57Bl/6 background. In addition, due to the known differences in the cardiovascular systems of males and females, we identified important biological sex differences. One group compared the different rates of regression of heart weights between Iso and Ang II and showed that regression occurred from both stimuli^(20)^. However, the timing and dosage of these experiments differ from our current study^(20)^. Further, no potential mechanisms in this previous study were investigated, and ours is the first to compare males and females in either of these models.

The findings reported here are summarized in Figure 7. There were no significant differences in the hypertrophic responses to either agonist or sex. However, with respect to regression, there were both sex differences and agonist-specific differences (Figure 1). After withdrawal of Iso treatment, males completely regressed by 4 days, whereas it took females 7 days to completely regress (Figure 1). In response to Ang II withdrawal, males did not experience any significant regression and females had significant regression at P4. However, LV mass was still significantly larger than the vehicle control after 7 days in both males and females (Figure 1). A previous study showed that Ang II produces a significant amount of fibrosis ^(29)^, which could be an underlying cause of irreversible hypertrophy. While Ang II agonism in one study showed complete regression after 7 days, the dose was lower in this study and the mice did not experience as much hypertrophy^(20)^. We have shown for the first time significant differences in the rate of regression comparing males and females.

**Figure 7:**
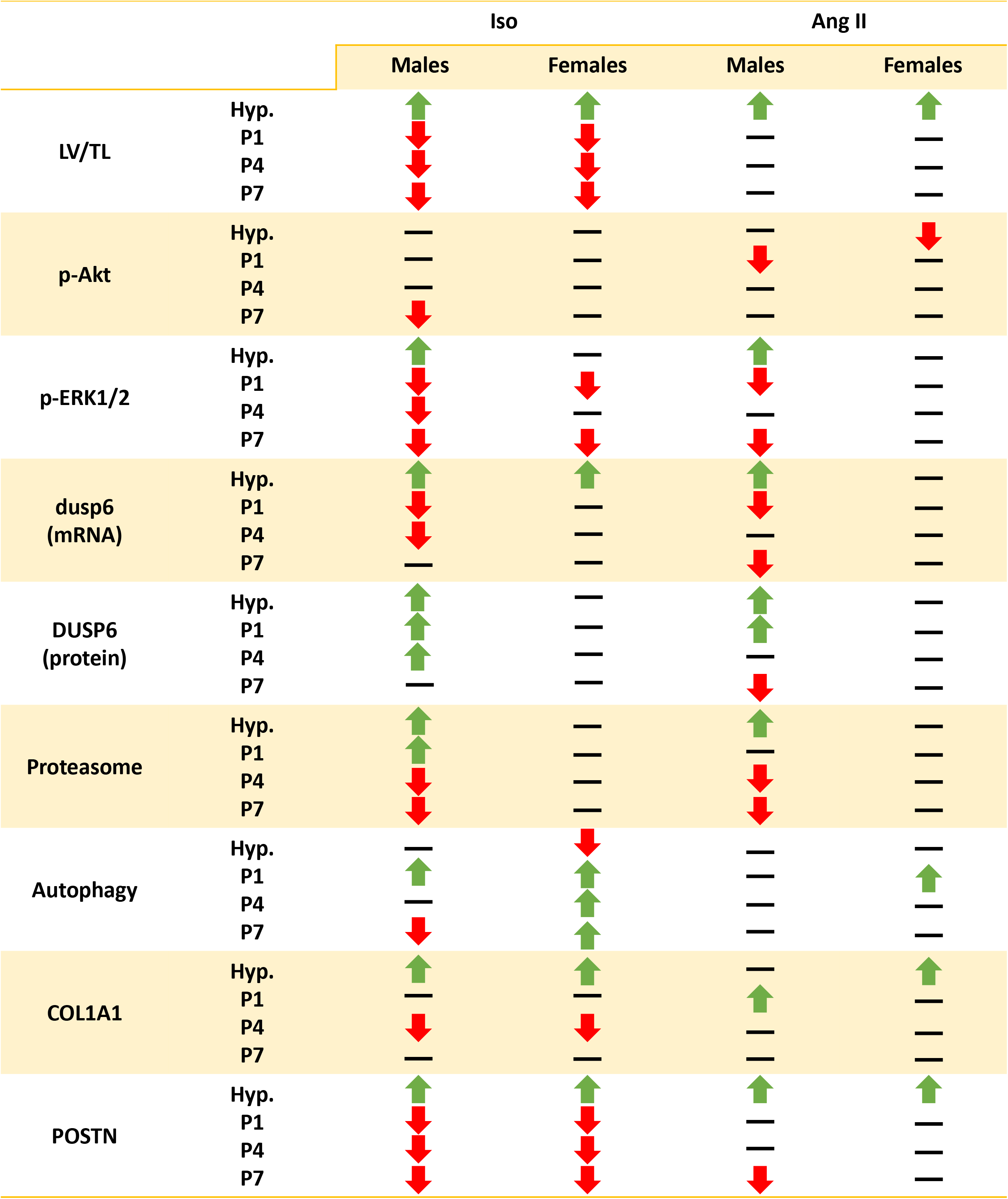
Conclusions table. Hypertrophy groups are compared to vehicle controls and regression groups are compared to the hypertrophy group.

We assessed the phosphorylation status of a number of signaling molecules. There were no significant changes in p-Akt levels during Iso or Ang II induced hypertrophy in males. However, there was a decrease in p-Akt at P7 in Iso treated male mice compared to the hypertrophic timepoint (Supplemental Figure 2A). In addition, there was a significant increase in p-ERK1/2 during Iso or Ang II induced hypertrophy in males followed by a decrease immediately after the removal of either agonist (Figure 3A). Activation of ERK1/2 has been observed in many pathological models^(42)^ and inhibition of p-ERK1/2 has been shown to inhibit cardiac hypertrophy^(43, 44)^. However, ours is the first report to show the rapid inactivation of ERK1/2 following the removal of a pathological stimulus. In contrast to male mice, female mice showed very minimal responses in these signaling pathways during hypertrophy, with only a significant decrease of p-Akt in response to Ang II (Supplemental Figure 2B). During regression, p-ERK1/2 and p-Akt were significantly decreased following the removal of Iso, and p-Akt was significantly decreased following Ang II removal (Figure 3C & Supplemental Figure 2B). These minimal changes during hypertrophy could be due to the higher level of active kinases in females at baseline (Figure 2), and therefore the response was not needed because levels were already high enough, or the threshold of the kinase had already been reached. These sex differences in kinases observed at baseline are similar to previously reported studies both in humans and mice ^(45)^.

As a follow-up to the increased ERK1/2 activity in males due to Iso and Ang II, DUSP6 was measured as a potential regulator. mRNA levels were elevated in response to Iso and Ang II in males and in response to Ang II in females (Figure 3D, E). Dusp6 mRNA may be responding to the activation of ERK1/2 during the hypertrophic response and is primed to be translated when p-ERK1/2 is no longer required. This is shown at the first regression time-point when mRNA levels were significantly reduced at the same time-point when ERK1/2 is no longer active. DUSP6 protein remained high in males throughout Iso-induced regression to possibly maintain low levels of p-ERK1/2 and prevent its reactivation (Figure 3G). In females, the elevated levels of dusp6 mRNA may be a response to Iso; however, it appeared the mRNA was not translated as there were no significant changes in DUSP6 protein levels. DUSP6 protein levels were significantly increased in male mice in response to Ang II. At P1, DUSP6 protein remained elevated in males and was significantly increased in females. By P4, DUSP6 returned to baseline levels in the male mice. However, levels remained significantly elevated in female mice at P4 and P7.

Due to an increase in protein synthesis during hypertrophy, we hypothesized protein degradation pathways would have a significant role in regression of hypertrophy. First, we found that proteasome activity was increased with Iso and further increased immediately after removal of Iso in males (Figure 4B). In contrast, Ang II resulted in a greater increase in proteasome activity compared to Iso; however, the activity level was significantly reduced immediately following the withdrawal of Ang II (P1). These results indicate the proteasome may have an active role in regression from Iso induced hypertrophy, but not following the removal of Ang II, in male mice. There were no notable changes in proteasome activity in female mice (Iso or Ang II) (Figure 4C), nor were there any sex differences observed at baseline (Figure 4A). To assess whether autophagy was playing a role in the regression of hypertrophy, we measured the ratio of LC3II to LC3I, and found that females had an increased level of autophagy compared to males at baseline (Figure 4D). Some reports indicate autophagy can be an indicator of health, as autophagy tends to decrease with age and poor health ^(46)^. Further, when autophagy was promoted by caloric restriction, diastolic dysfunction was delayed in an aging rodent^(47)^ whereas when autophagy was inhibited, cardiac function and structure declined ^(46)^. This could be one explanation as to why females respond better to some cardiac stresses. Reports vary on the autophagic response to pathological hypertrophy, with some reporting autophagy was increased ^(48)^ while others reported autophagy was decreased during early hypertrophy^(49, 50)^, and then increased in failing hearts ^(50)^. However, each of these studies only reported on male rodents, and because autophagy is an important target for many pharmaceutical treatments, it is important to understand how autophagy is differently regulated in males and females in response to cardiac stresses. We showed in this report autophagy was decreased, but only in females treated with Iso (Figure 4F). Iso did not induce autophagy changes in male mice, or with Ang II in either sex. During regression, the level of autophagy increased in both males and females compared to the Iso hypertrophic state; however, the increase was greater in males (Figure 4E, F). Because both autophagy and the proteasome were more active in males in response to the removal of Iso, this could be why regression occurs faster, and we believe this could be a determining factor in regulating regression of cardiac hypertrophy. Removal of Ang II also elicited an increase in autophagy, but only in females.

Regression of cardiac hypertrophy could be hindered by the deposition of collagen, and the ability of the heart to degrade a fibrotic network. While fibrosis was not associated with Iso in this mouse model ^(40, 41)^, interstitial fibrosis did occur with Ang II ^(29, 38)^. Therefore, we were interested in investigating the activation of fibrotic genes, and whether the activation of these genes was able to be reversed. We saw an activation of fibrotic genes (COL1A1 and POSTN), with both Iso and Ang II, and in both sexes (Figure 5). However, there were differences in the amount of activation and how long the expression of these genes was elevated. Overall, Ang II induced a greater pathological induction of these ECM genes. It was then hypothesized, this could lead to increased collagen content, and if the collagen is unable to be degraded, this could be a leading cause of irreversibility. We found there was not an immediate increase in collagen content after 7 days of Ang II; however, following the removal of Ang II, there were significant increases (Figure 6B). Further, these increases correlated with the increases in ECM gene expression after Ang II removal, which did not occur during Iso induced regression. In addition, the collagen content did not decrease after withdrawal of Ang II. These results indicate a collagen network was possibly forming in the left ventricle and this was likely due to the prolonged expression of ECM genes, even after Ang II had been removed.

In conclusion, complete regression of pathological cardiac hypertrophy occurred in the Iso model in both sexes, but the rate of regression was slower in females. Incomplete regression occurred in the Ang II model. These differences could be due to alterations in signaling pathways, protein degradation pathways, or the accumulation of ECM proteins. Further, there were many sex differences associated with these two hypertrophic models. Future studies will include probing more pathways to further understand regression, especially any sex differences that may affect how patients are treated and respond to anti-hypertensive treatments.

## Acknowledgements

We would like to thank Dr. Claudia Crocini and Dr. Angela Peter for helpful discussions.

The authors have no conflicts of interest.

---

**Table 1:** Body and heart weights of Iso induced hypertrophy and regression mice. P Post-Stimulus Day; BW Body Weight; TL Tibia Length; HW Heart Weight; LV Left Ventricle; RV Right Ventricle. One-way ANOVA Post hoc-Uncorrected Fisher’s LSD. *p<.05; **p<.01; ***p<.001; ****p<.0001 significance from vehicle control.

|  | Males |  |  |  |  | Females |  |  |  |  |
| --- | --- | --- | --- | --- | --- | --- | --- | --- | --- | --- |
|  | Veh | Iso | P1 | P4 | P7 | Veh | Iso | P1 | P4 | P7 |
| **BW/TL**<br>g/mm | 1.61 | 1.67 | 1.66 | 1.68 | 1.66 | 1.33 | 1.35 | 1.34 | 1.37 | 1.36 |
| **HW/TL**<br>mg/mm | 6.83 | 9.39<br>**** | 7.96<br>**** | 7.56 | 7.36 | 5.87 | 7.66<br>**** | 6.42<br>**** | 6.38<br>** | 5.97 |
| **LV/TL**<br>mg/mm | 4.97 | 6.65<br>**** | 5.77<br>** | 5.56 | 5.39 | 4.27 | 5.49<br>**** | 4.70<br>** | 4.68<br>* | 4.34 |
| **RV/TL**<br>mg/mm | 1.19 | 1.57<br>**** | 1.29 | 1.23 | 1.19 | 0.94 | 1.22<br>**** | 1.05<br>** | 1.01 | 1.02 |

**Table 2:** Body and heart weights of Ang II induced hypertrophy and regression mice. P Post-Stimulus Day; BW Body Weight; TL Tibia Length; HW Heart Weight; LV Left Ventricle; RV Right Ventricle. One-way ANOVA Post hoc-Uncorrected Fisher’s LSD. +p<.05; ++p<.01; +++p<.001; ++++p<.0001 significance from vehicle control.

|  | Males |  |  |  |  | Females |  |  |  |  |
| --- | --- | --- | --- | --- | --- | --- | --- | --- | --- | --- |
|  | Veh | Ang II | P1 | P4 | P7 | Veh | Ang II | P1 | P4 | P7 |
| **BW/TL**<br>g/mm | 1.62 | 1.42 | 1.42 | 1.43 | 1.47 | 1.39 | 1.36 | 1.27 | 1.26 | 1.31 |
| **HW/TL**<br>mg/mm | 6.73 | 8.00<br>++ | 7.81<br>++ | 7.91<br>+ | 7.73<br>++ | 5.71 | 7.38<br>++++ | 7.25<br>+++ | 6.78<br>+ | 6.91<br>++ |
| **LV/TL**<br>mg/mm | 4.70 | 6.02<br>+++ | 5.78<br>++ | 5.64<br>+ | 5.69<br>+ | 4.20 | 5.65<br>++++ | 5.40<br>+++ | 5.07<br>++ | 5.03<br>++ |
| **RV/TL**<br>mg/mm | 1.18 | 1.14 | 1.14 | 1.24 | 1.24 | 0.93 | 0.96 | 0.98 | 1.15<br>+ | 1.07 |

