## Supplemental Figures for "Regression from pathological hypertrophy is sexually dimorphic and stimulus-specific"

### Supplemental: Phospho-kinase array

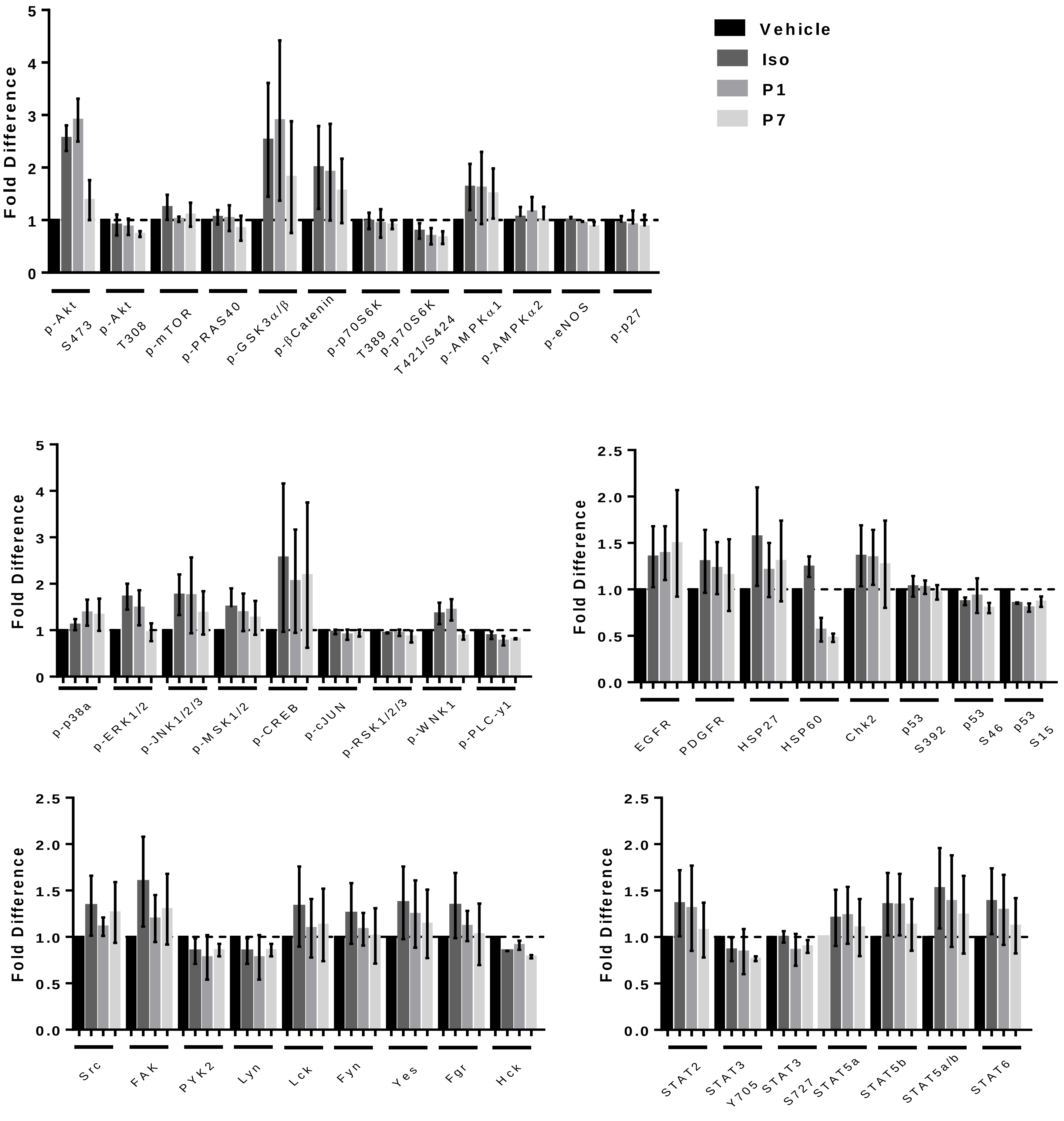


**Supplemental Figure 1: Phospho-kinase array blot quantified.** Male mice. n=2. Mean ± SEM.

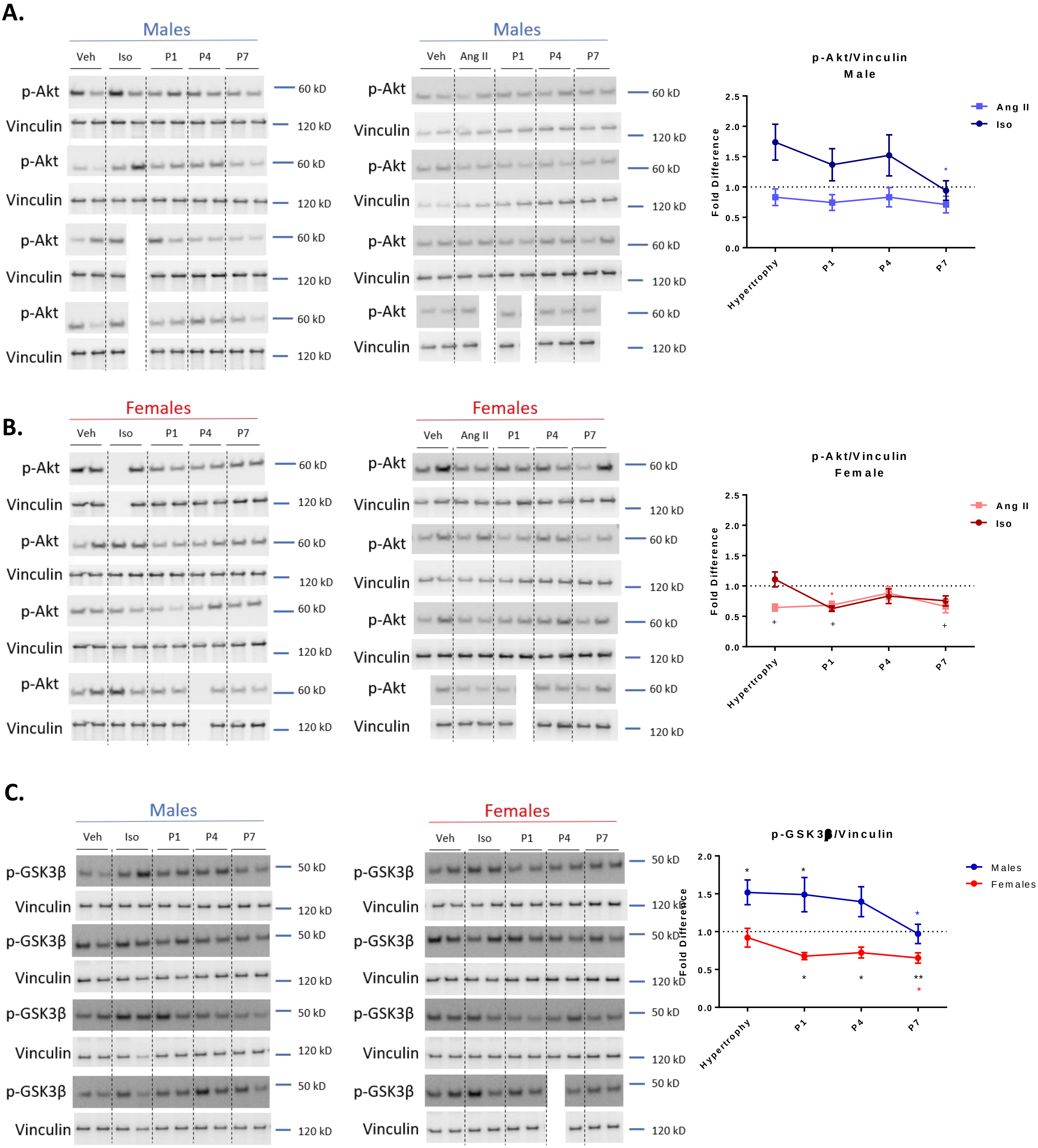

A.
Males
Veh
Iso
P1
P4
P7
p-Akt
Vinculin
60 kD
120 kD
Males
Veh
Ang II
P1
P4
P7
p-Akt/Vinculin
Male
Ang II
Iso
Fold Difference
2.5
2.0
1.5
1.0
0.5
0.0
Hypertrophy
P1
P4
P7
B.
Females
Veh
Iso
P1
P4
P7
p-Akt
Vinculin
60 kD
120 kD
Females
Veh
Ang II
P1
P4
P7
p-Akt/Vinculin
Female
Ang II
Iso
Fold Difference
Hypertrophy
P1
P4
P7
C.
Males
Veh
Iso
P1
P4
P7
p-GSK3β
Vinculin
50 kD
120 kD
Females
Veh
Iso
P1
P4
P7
p-GSK3β/Vinculin
Males
Females
Fold Difference
2.0
1.5
1.0
0.5
0.0
Hypertrophy
P1
P4
P7

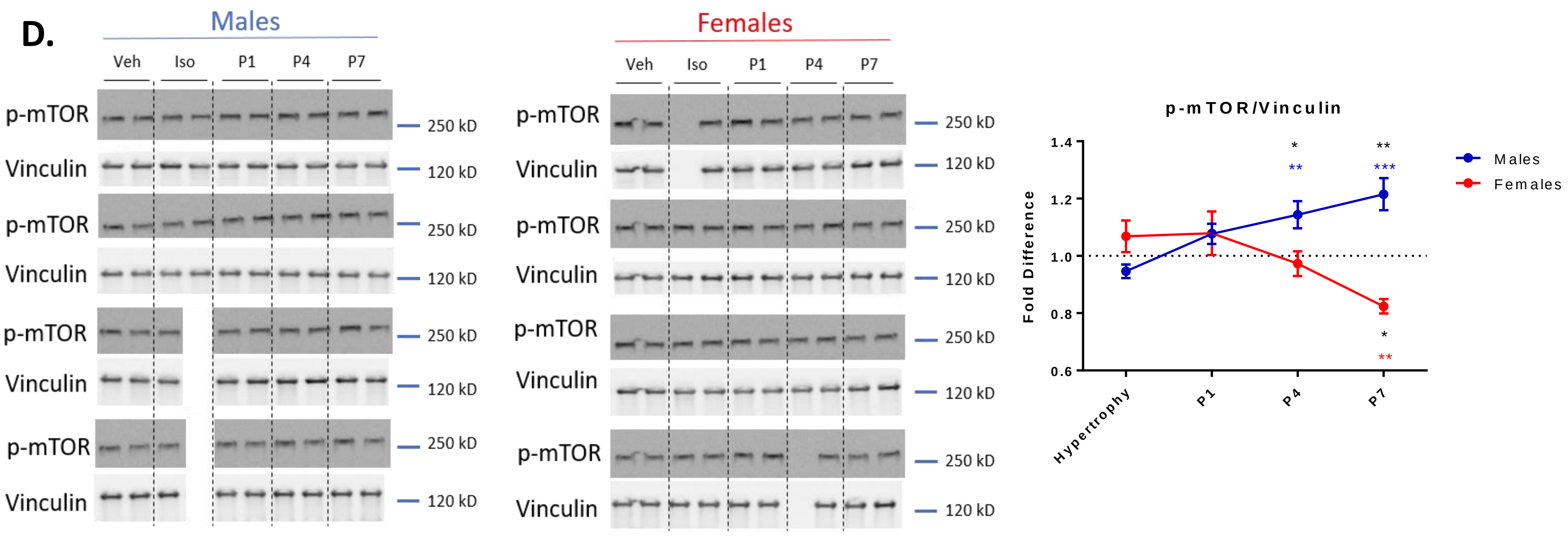


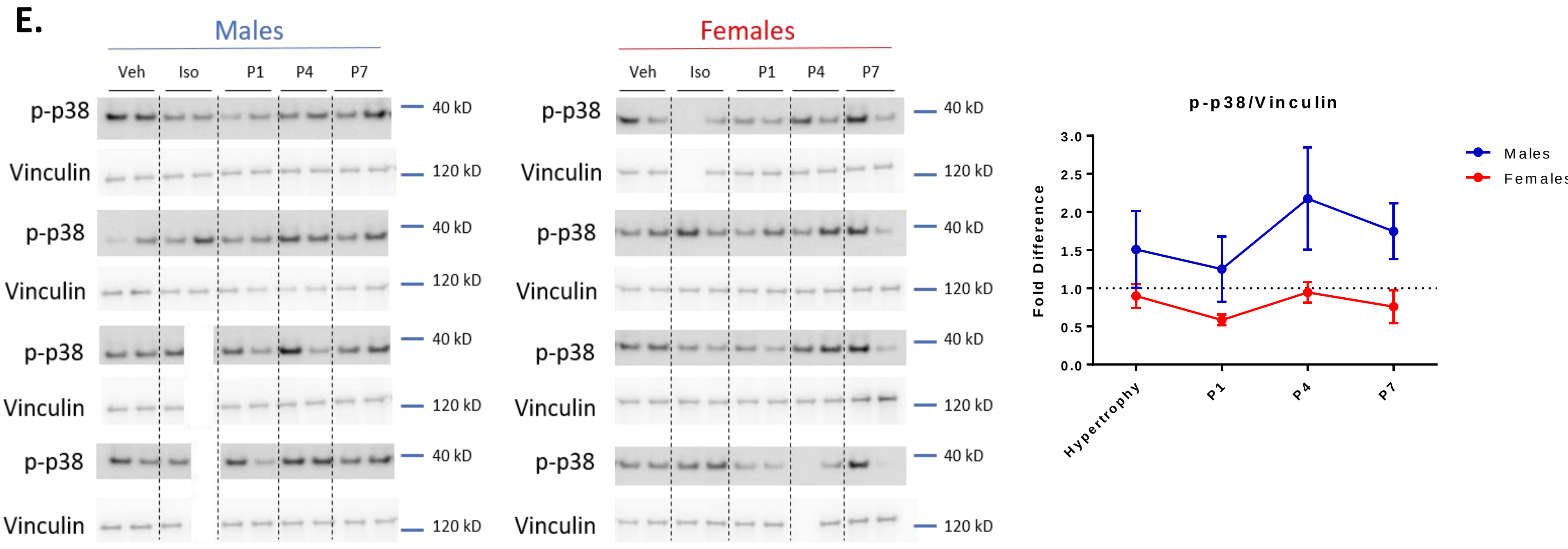


**Supplemental 2: Signaling pathways during Iso hypertrophy & regression.** A. Akt does not respond to Iso- or Ang II-induced hypertrophy or following the removal of either stimulus in males B. Akt does not respond to Iso- or Ang II-induced hypertrophy or following the removal of either stimulus in females C. GSK3β is activated in response Iso then decreases after 4 days of Iso removal in males. GSK3β is inactivated in response to Iso removal in females D. mTOR is increased in response to Iso removal in males and decreased in response to Iso removal in females E. p38 is not altered in response to Iso in either males or females. Values are normalized to Vinculin. n=6-8/group. Mean ±SEM One-way ANOVA Post hoc-Uncorrected Fisher's LSD. *p<.05 * significance in Iso group. + significance in Ang II group. */+significance from vehicle control. */+; */+ significance from hypertrophy.

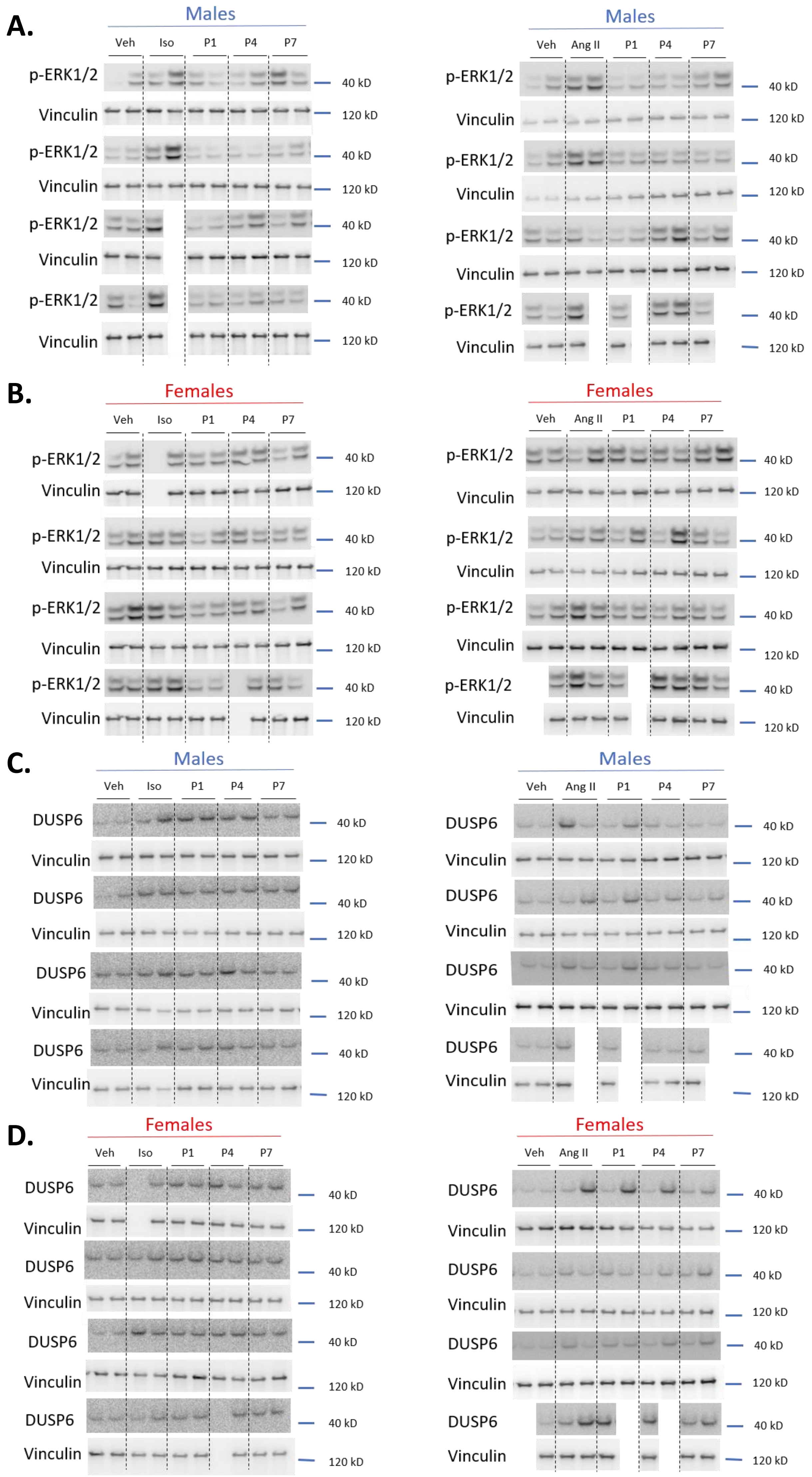


**Supplemental Figure 3: Western blots of p-ERK1/2 and DUSP6 in response to hypertrophy & regression** A. p-ERK1/2 in male mice B. p-ERK1/2 in female mice. C. DUSP6 protein in male mice. D. DUSP6 protein in female mice. Protein normalized to Vinculin. n=6-8/group.

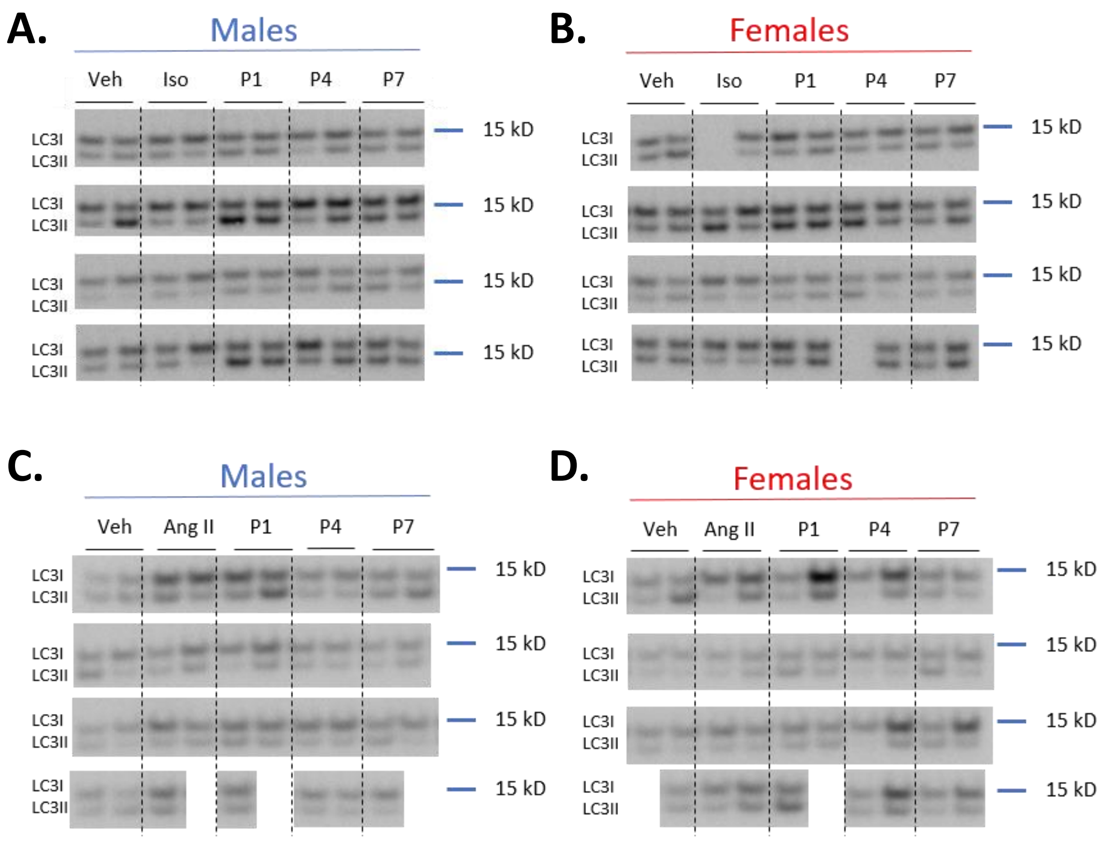


**Supplemental 4: LC3 Western blots:** A. LC3 in Iso treated male mice B. LC3 in Iso treated female mice. C. LC3 in Ang II treated male mice. D. LC3 in Ang II treated female mice
